# Dopaminergic amacrine cells express HCN channels in the developing and adult mouse retina

**DOI:** 10.1101/2024.07.20.604440

**Authors:** Emilio J Romano, Dao-Qi Zhang

**Affiliations:** Eye Research Institute, Oakland University, Rochester, Michigan; Eye Research Center, Oakland University William Beaumont School of Medicine, Rochester, Michigan

## Abstract

**Purpose:** To determine the molecular and functional expression of hyperpolarization-activated cyclic nucleotide-gated (HCN) channels in developing and mature dopaminergic amacrine cells (DACs), the sole source of ocular dopamine that plays a vital role in visual function and eye development.

**Methods:** HCN channels are encoded by isoforms 1-4. HCN1, HCN2, and HCN4 were immunostained in retinal slices obtained from mice at postnatal day 4 (P4), P8, and P12 as well as in adults. Each HCN channel isoform was also immunostained with tyrosine hydroxylase, a marker for DACs, at P12 and adult retinas. Genetically-marked DACs were recorded in flat-mount retina preparation using a whole-cell current-clamp technique.

**Results:** HCN1 was expressed in rods/cones, amacrine cells, and retinal ganglion cells (RGCs) at P4, along with bipolar cells by P12. Different from HCN1, HCN2 and HCN4 were each expressed in amacrine cells and RGCs at P4, along with bipolar cells by P8, and in rods/cones by P12. Double immunostaining shows that each of the three isoforms was expressed in approximately half of DACs at P12 but in almost all DACs in adults. Electrophysiology results demonstrate that HCN channel isoforms form functional HCN channels, and the pharmacological blockade of HCN channels reduced the spontaneous firing frequency in most DACs.

**Conclusions:** Each class of retinal neurons may use different isoforms of HCN channels to function during development. HCN1, HCN2, and HCN4 form functional HCN channels in DACs, which appears to modulate their spontaneous firing activity.

## Introduction

Visual signals generated by rod and cone photoreceptors are transmitted from bipolar cells to retinal ganglion cells (RGCs) and then, through the optic nerve, to the brain’s visual centers, generating vision. The signal transmission through this neural pathway is shaped by neuromodulators, including dopamine^1^. The sole source of retinal dopamine is dopaminergic amacrine cells (DACs), a subclass of inhibitory amacrine cells. DACs are also called interplexiform cells due to their processes stratifying in both the outer plexiform layer (OPL) and inner plexiform layer (IPL). Therefore, dopamine released from DACs can be diffused throughout the retina, regulating circadian rhythm within the eye and reconfiguring neural circuits for visual information processing^2-7^.

DACs can generate action potentials that trigger dopamine release. Cultured DACs can spontaneously fire action potentials in a rhythmic pattern through the activation of persistent sodium channels^8^. Ex vivo DACs also spontaneously fire action potentials but in an arrhythmic pattern with a mixture of single and bursting activity^9^. However, the ionic mechanisms underlying arrhythmic spontaneous action potential generation remain poorly understood. Hyperpolarization-activated cyclic nucleotide-gated channels (HCN), encoded by isoforms 1-4, play a key role in controlling the firing of neurons in the central nervous system. A line of evidence has shown that HCN channels are expressed in rod and cone photoreceptors^10-15^, bipolar cells^16-20^, and RGCs^21-24^, which modulate excitatory signal transmission from the outer retina to the inner retina. Both HCN1 and HCN2 channels are also present in amacrine cells of the retina^16, 23, 25^, which may be involved in lateral inhibition processing. However, the specific type of amacrine cells that HCN channels are located on has not been identified. Therefore, it is still unclear whether HCN channels are expressed on DACs, regulating their spontaneous firing activity.

DACs appear almost immediately after birth in mice and dopamine release from them is observed prior to visual experience, contributing to retinal neural circuit development^26, 27^. Therefore, it would be interesting to determine whether HCN channels are expressed in both developing and mature DACs, regulating their spontaneous activity. Doing this requires exploring the unknown expression and distribution of HCN channel subunits throughout the developing retina. Understanding the development of HCN channels would provide insights into the mechanisms by which HCN channels are involved in the development of the visual system in general. Particularly, HCN channels in cone bipolar cells contribute to the generation of glutamatergic spontaneous activity (termed glutamatergic retinal waves) in the second postnatal week of mice^20^. In addition, HCN channels can mediate phototransduction in intrinsically photosensitive RGCs (ipRGCs) which plays a critical role in neural circuit development before visual experience^22, 24, 28^.

In the present study, we first detected the expression of HCN channel proteins in the developing retina. We then determined the expression of each HCN channel isoform on DACs in developing and adult retinas. Finally, we examined functional HCN channels on DACs and their role in DAC spontaneous activity generation.

## Materials and Methods

### Animals

Wild-type C57BL/6 mice were obtained from the Jackson Laboratory (Bar Harbor, ME, USA). Transgenic mice in which DACs are genetically labeled with a red fluorescent protein (RFP) under the rat *tyrosine hydroxylase* (*TH*) promoter were created on a C57BL/6J background at Vanderbilt University^29^. All mice used in this study were housed at Oakland University’s Biomedical Research Support Facility in a 12/12-hour light/dark cycle. All animal care and procedures were completed in line with the guidelines from the National Institutes of Health and the ARVO Animal Statement for the Use of Animals in Ophthalmic and Vision Research. The use of the animals was approved by the Institutional Animal Care and Use Committee at Oakland University.

### Immunohistochemistry

Mice were sacrificed by an overdose of carbon dioxide followed by cervical dislocation. Eyeballs from mice were enucleated, fixed, and sliced following the procedures described previously^30, 31^. Vertical retinal slices were incubated with 1% BSA and 0.3% Triton-X 100 for two hours at room temperature. Following this, retinal slices were incubated in primary antibodies either overnight at room temperature or over two nights at 4°C. Primary antibodies used in this study were against HCN1, HCN2, HCN4 (all raised in rabbit; used at a concentration of 1:1000; Alamone Labs, Jerusalem, Israel), and TH (raised in sheep; used at a concentration of 1:2000; EMD Millipore, Burlington, MA, USA). Following incubation in primary antibodies, retinal slices were rinsed using 0.1X PBS and then incubated with secondary antibodies in blocking solution for two hours at room temperature. Secondary antibodies were raised in donkey and were conjugated to AlexaFluor-488 or 594 (used at a 1:500 concentration; Life Technologies, Carlsbad, California, USA). Following incubation in secondary antibodies, retinal slices were rinsed again using PBS and then mounted using Vectashield Hard-Set mounting solution (Vector Laboratories, Burlingame, CA, USA).

### Imaging and Analysis

Vertical retinal slices were visualized on a ZEISS LSM 900 with Airyscan 2 Confocal Microscope using Zeiss Zen software to take images (Zeiss, Oberkochen, Germany). Images were taken using 20X or 40X magnification. Image analysis and processing were also done using Zeiss Zen software.

### Electrophysiology

Following euthanasia, eyeballs were enucleated under dim red illumination and transferred to a petri dish filled with oxygenated extracellular solution containing Ames’ medium (MyBioSource, San Diego, CA). Under dim red light, the cornea and lens were removed from the eyeballs, and the retina was separated from the sclera. The retina was then placed with the photoreceptor side down in a recording chamber mounted on the stage of an upright conventional fluorescence microscope (BX51WI, Olympus, Tokyo, Japan). Oxygenated Ames’ medium (pH 7.4 bubbled with 95% O_2_-5% CO_2_) continuously perfused the recording chamber at a rate of 2–3 ml/min, and the superfusate was maintained at 32–34°C by a temperature control unit (TC-344B, Warner Instruments, Hamden, CT).

Cells and recording pipettes were viewed on a computer monitor coupled to a digital camera (XM10, Olympus, Tokyo, Japan) mounted on the microscope. *TH-*RFP expressing cells were randomly selected throughout the retina after being visualized by fluorescence using a rhodamine filter set. A peak wavelength of 535 nm from a fluorescence LED illumination system (pE-2, CoolLED Ltd., Andover, UK) was used to give a brief “snap-shot” of fluorescence excitation light (1–2 s).

Whole-cell current-clamp recordings were made from the soma of RFP-labelled DACs with 4–8 MΩ electrodes, and signals were amplified with an Axopatch 200B amplifier (Molecular Devices, Sunnyvale, CA). The intracellular solution for the whole cell current-clamp experiments contained (in mM) 120 K-gluconate, 2 EGTA, 10 HEPES, 4 KCl, 5 NaCl, 4 Na-ATP, and 0.3 Na-GTP. The liquid junction potential was 10.5 mV, which was corrected offline. All electrophysiological data were analyzed using Clampfit software (Molecular Devices, Sunnyvale, CA).

Statistical comparisons were made between them using the paired *t*-test. A student’s *t*-test was used for comparison between two independent groups. Values are presented as the mean ± SEM in the present study, and *p* < 0.05 was considered statistically significant.

## Results

### The expression of HCN channel isoforms in the developing retina

A line of evidence has shown that all four HCN channel isoforms are expressed in the adult retina ^16, 23^. However, the expression and distribution of HCN channels in the developing retina remain unknown. We immunostained HCN1, HCN2, and HCN4 on the retinas of mice at P4, P8, and P12. For comparison, similar experiments were also carried out in adult retinas. Consistent with adult retinas (Figure 1A)^16, 23^, HCN1 was expressed in the outer nuclear layer (ONL, rods, and cones), proximal part of the inner nuclear layer (INL, amacrine cells), IPL, and ganglion cell layer (GCL, amacrine cells, and RGCs) at P4, P8, and P12 (Figure 1A). Interestingly, HCN1 was barely detected in the distal part of the INL where bipolar cells are located until P12 (Figure 1A). Our data suggest that the expression of HCN1 appears in bipolar cells later than in rods/cones, amacrine cells, and RGCs.

**Figure 1.**
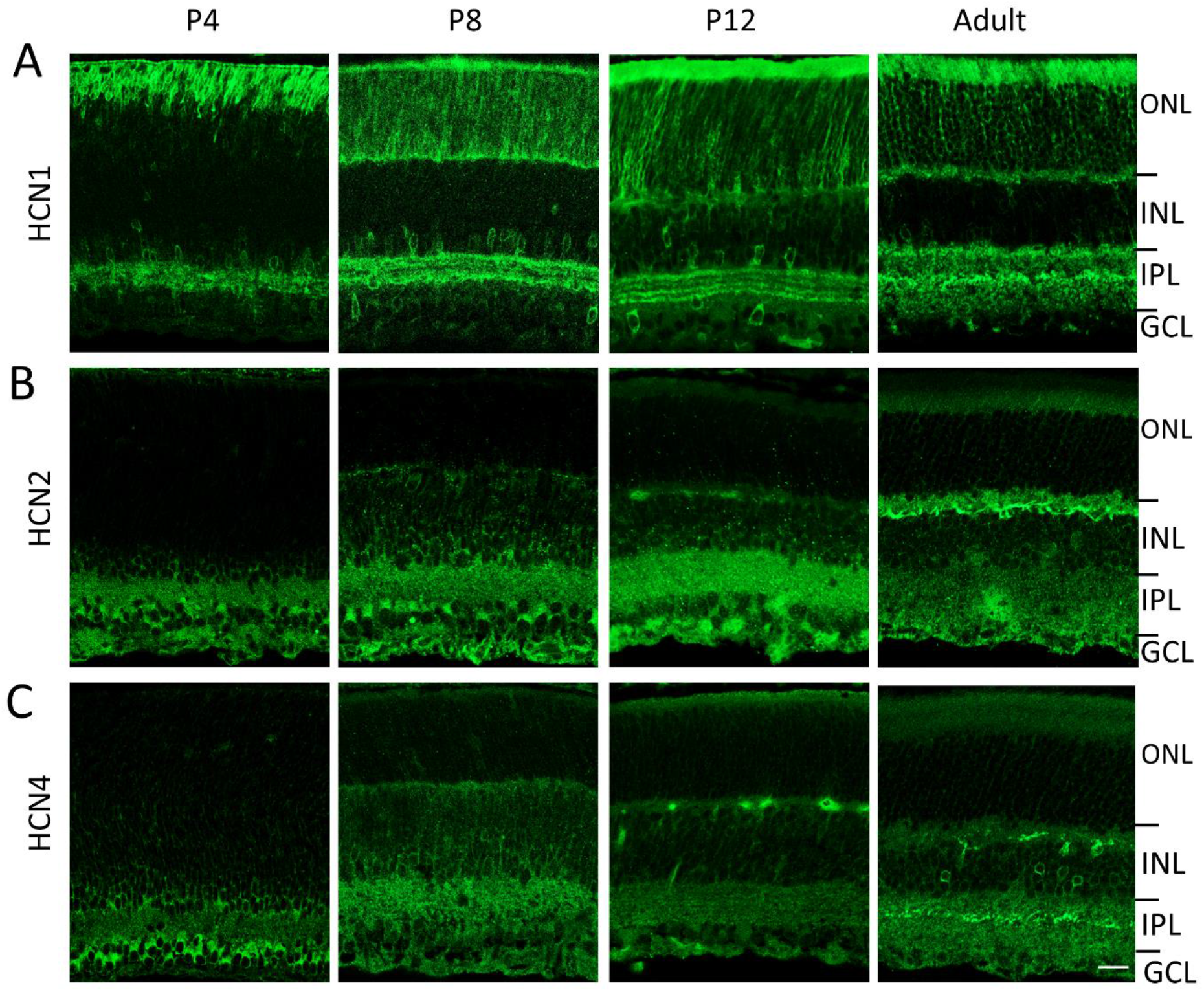
Differential expression of HCN channel isoforms on retinal neurons during the course of development. Vertical retinal slices were immunostained (green) with antibodies specific for HCN1 (**A**), HCN2 (**B**), and HCN4 (**C**). Retinal cell layers and plexiform layers are marked on the right side of the figure: ONL, outer nuclear layer; INL, inner nuclear layer; IPL, inner plexiform layer; GCL, ganglion cell layer. **A**. HCN 1 was expressed in the ONL, inner part of INL, and GCL at P4 (far left panel) and P8 (middle left panel). Additional expression was observed in the outer part of the INL at P12 (middle right panel) and in adults (far right panel). **B**. HCN2 expression began in the inner part of the INL and GCL at P4 (far left panel) and then extended to the outer part of the INL at P8 (middle left panel). Additional expression was observed in the ONL at P12 (middle right panel) and in adults (far right panel). **C**. The distribution patterns of HCN4 expression are similar to those of HCN2 expression shown in B. Scar bar, 20 µm.

Similar to HCN1, HCN2 (Figure 1B) and HCN4 (Figure 1C) each began to be expressed in the proximal part of the INL, IPL, and GCL at P4. However, different from HCN1, HCN2 (Figure 1B) and HCN4 (Figure 1C) each began to appear in the distal part of the INL at P8 and in the ONL until P12. The results suggest that HCN2 and HCN4 appear in amacrine cells and RGCs earlier than in photoreceptors and bipolar cells.

We also attempted to detect the expression of HCN3 in developing and adult retina. However, the two antibodies we used to agonist HCN3 did not work well on retinal tissue. Therefore, we do not include the data in this work.

### HCN channel isoforms are expressed in a subpopulation of DACs during development

To determine whether developing DACs express HCN channels, we performed double immunostaining of TH with each one of the HCN channel antibodies on vertical retina slices obtained from mice at P12, the age in which all three HCN channel isoforms were expressed in all classes of retinal neurons (Figure 1). Double labeling with the TH antibody shows that the HCN1 channel isoform was expressed on the membrane of DACs. The expression on DACs (white arrow) was relatively weak comparing that on bipolar cells (asterisk) in the INL (Figure 2A). Interestingly, HCN1 expression was not observed in every tested DAC. We counted the number of DACs from the vertical retinal slices obtained from four pups. Of 24 DACs counted, ∼46% of them expressed HCN1 channel isoform (Figure 2A). DACs also contained HCN2 (Figure 2B) and HCN4 (Figure 2C) channel isoforms. The percentage of HCN2-containing DACs (18 out of 26 cells) was higher than that of HCN1-containing DACs (∼69% vs. ∼46%) (Figure 2B). The percentage of HCN4-containing DACs (12 out of 16 cells) was also higher than that of HCN1-containing DACs (75% vs ∼46%) as well as than that of HCN2-containing DACs (75% vs ∼69%) (Figure 2C). The results suggest that each of the HCN channel isoforms studied (HCN1, HCN2, and HCN4) are expressed in the majority of DACs around eye-opening.

**Figure 2.**
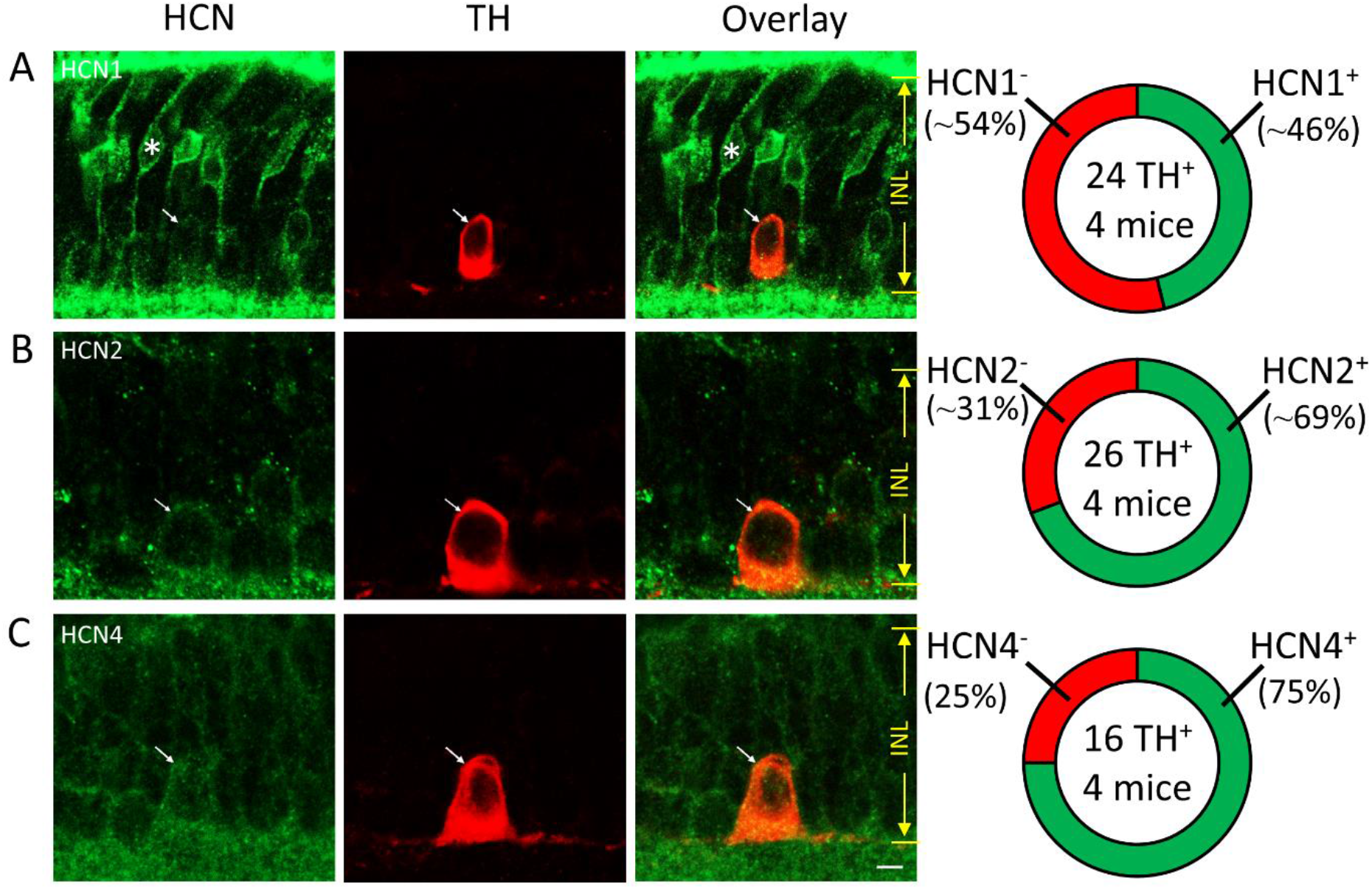
Developing DACs express HCN1, HCN2, and HCN4 channel subunits. Double immunostaining of TH with HCN1 (A), HCN2 (B), or HCN4 (C) in vertical retinal slices obtained from mice at P12. Images in the far-left panel demonstrate immunostaining for each of the three HCN channel isoforms (green). TH-containing DACs (middle-left panel) expressed HCN1 (A), HCN2 (B), or HCN4 (C) (overlay images in the middle-right panel). White arrows indicate DACs. The white asterisk indicates a bipolar cell. Pie charts on the far-right panel show the percentages of DACs negative and positive to HCN1 (A), HCN2 (B), or HCN4 (C). The total number of DACs illustrated in the center of each chart were counted in retinal vertical slices obtained from 4 mice. Scale bar: 5 µm.

### HCN channel isoforms are expressed in almost all mature DACs

One possibility for not detecting one of the three HCN isoforms in every DAC is that HCN channels are not fully developed in them before eye-opening. To test this possibility, we determined the expression of HCN channels of DACs in the adult retina. Figure 3 illustrates the co-localization of TH with either HCN1 (Figure 3A), HCN2 (Figure 3B), or HCN4 (Figure 3C). We counted the co-labeled DACs from vertical slices obtained from three adult mice. We found that 95% of DACs (38 out 40 cells) examined expressed HCN1 (Figure 3A) and 100% of DACs examined expressed HCN2 (28 out of 28 cells, Figure 3B) or HCN4 (22 out of 22 cells, Figure 3C). Our data suggest that DACs co-express all three isoforms of the tested HCN channels in the adult retina.

**Figure 3.**
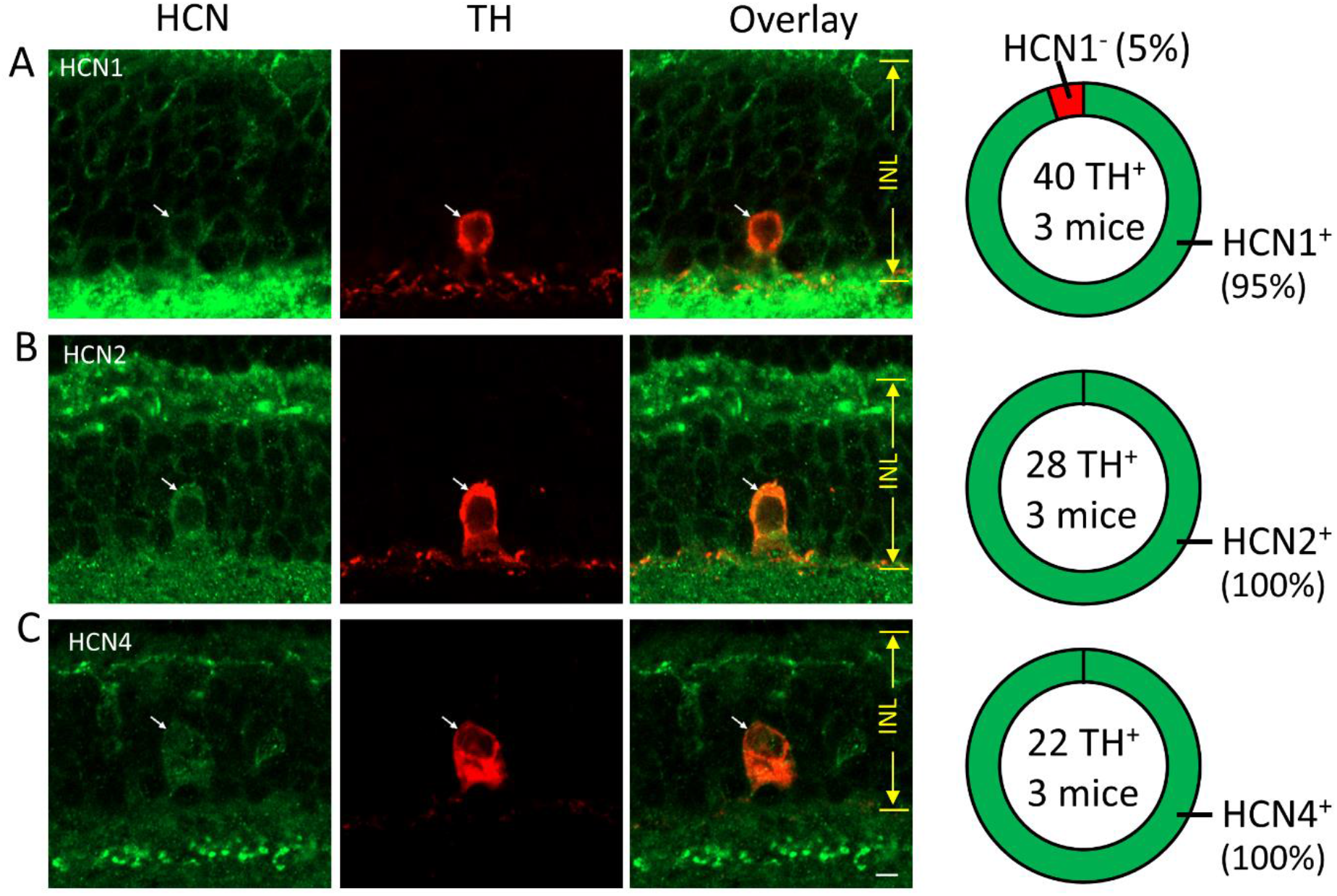
DACs express HCN1, HCN2, and HCN4 channel subunits in the adult retina. Double immunostaining of TH with HCN1 (**A**), HCN2 (**B**), or HCN4 (**C**) in vertical retinal slices obtained from adult mice. Images in the far-left panel demonstrate immunostaining for each of three HCN channel isoforms (green). TH positive DACs (middle-left panel) contained HCN1 (**A**), HCN2 (**B**), or HCN4 (**C**) (overlay images in the middle-right panel). Pie charts on the far-right panel show the percentages of DACs negative and positive to HCN1 (**A**), HCN2 (**B**), or HCN4 (**C**). The total number of DACs illustrated in the center of each chart was counted in retinal vertical slices obtained from 3 mice. Scale bar: 5 µm.

### The majority of mature DACs exhibit a voltage sag in response to hyperpolarization

HCN channels are activated by membrane hyperpolarization and are permeable to both Na^+^ and K^+^ ions. At hyperpolarized membrane potentials, a rebound depolarization (sag) is thought to be the signature of the functional expression of HCN channels in neurons ^32, 33^. To reveal the voltage-sag, we performed whole-cell current-clamp recordings from DACs. DACs were genetically labeled with RFP (Figure 4A); therefore, we visualized them in the flat-mount retina using fluorescence microscopy and targeted them for patch-clamp recording under DIC (Figure 4B). Once we obtained whole-cell current-clamp recordings, a series of negative currents from 0 pA to -100 pA with a step of 20 pA were injected into a DAC. The currents induced hyperpolarization of the cell. We found that 16 out of 19 DACs recorded exhibited hyperpolarization-induced voltage sag when the cells were hyperpolarized below -80 mV (sag-expressing cells, Figure 4C). The remaining three cells did not show voltage sage in response hyperpolarization (no-sag-expressing cells, Figure 4C). The results suggest that the majority of DACs potentially express functional HCN channels.

**Figure 4.**
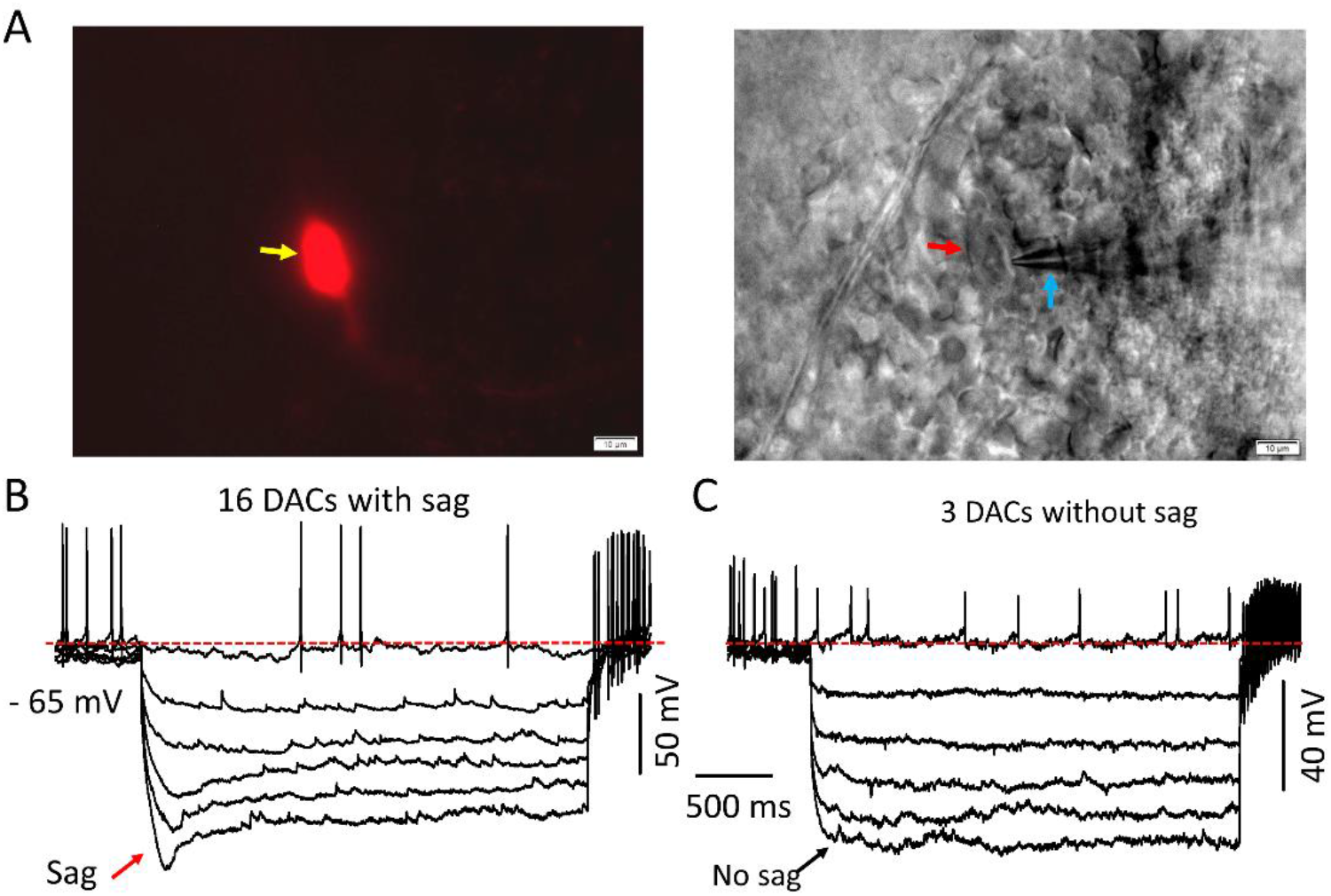
Biophysical property of functional HCN channels in DACs. A. A red fluorescent protein (RFP) DAC (left image) was visualized in an adult flat-mount retina under the differential interference contrast for patch-clamp recording (right image). **B**. Whole-cell current-clamp recordings were made on RFP-marked DACs. A series of negative currents from 0 to -100 pA with a step of 20 pA was injected into a cell, and they induced hyperpolarization of the cell. 16 out of 19 DACs recorded exhibited voltage sag when the cells were hyperpolarized below -80 mV. **C**. 3 out of 19 DACs did not show the voltage sag in response to hyperpolarization.

### HCN channel blockers reduce the amplitude of hyperpolarization-induced voltage-sag in DACs

To validate that DACs express functional channels, we used specific channel blockers. Cs^+^ and ZD7288 are two widely used blockers for HCN channels. The former in the millimolar range can effectively block HCN channels from the extracellular side by blocking the ion-conducting pore^34, 35^. In contrast, ZD7288 must diffuse through the cell membrane to bind to the intracellular vestibule of the ion-conducting pore to block HCN channels^36^. Figure 5A illustrates an example of the blockade of the voltage sag of a DAC by 2 mM Cs^+^. Averaged data show that Cs^+^ significantly reduced the amplitude of voltage sags at -80 mV was reduced from 17.39 ± 5.95 mV (mean ± S.E.) to 5.42 ± 2.31 mV (Figure 5B, P<0.05, n=6). Similar results were obtained using 30 µM ZD7288 (Figure 5C). ZD7288 decreased the voltage sag amplitude from 16.22 ± 3.26 mV to 6.88 ± 2.19 mV (Figure 5D, P<0.05, n=5). Overall, the results indicate that functional HCN channels are expressed on DACs.

**Figure 5.**
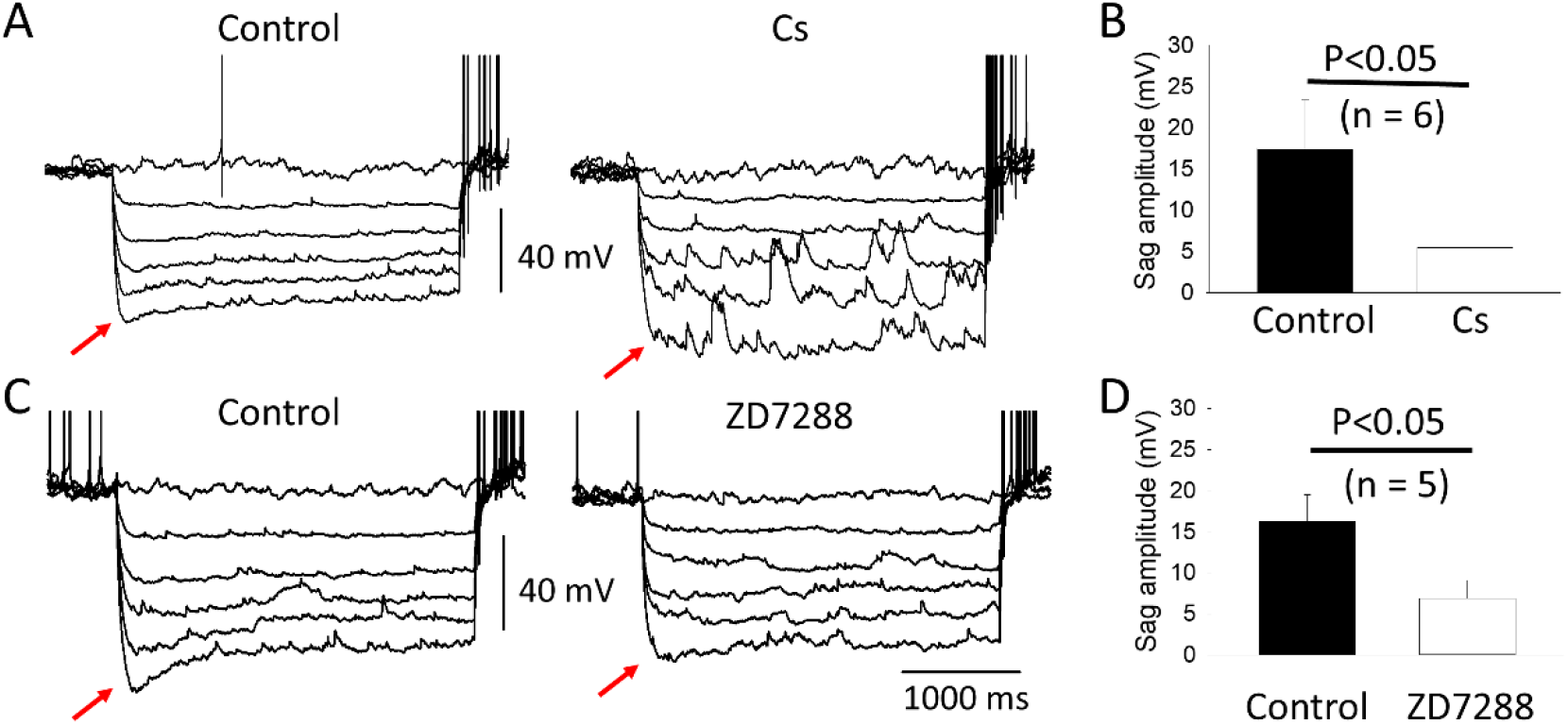
Pharmacological property of HCN channels in DACs. Whole-cell current-clamp recordings were made on RFP-marked DACs in flat-mount retinas as described in Figure 4. 2mM Cs^+^ (**A**) and 30 µM ZD7288 (**B**) reduced the peak amplitude of the voltage sag in DACs (indicated by red arrows), respectively. Mean data demonstrate that the reduction caused by Cs^+^ (**C**) and ZD7288 (**D**) was significant (P<0.05).

### Blockade of HCN channels reduces spontaneous activity in the majority of DACs

DACs exhibit spontaneous action potentials in darkness and the spontaneous activity is generated intrinsically^9^. To determine whether HCN channels are involved in the generation of spontaneous activity, we examined the effect of HCN channel blockers on the resting membrane potentials (RMPs) and spontaneous action potential frequency of DACs. Our results demonstrate that neither Cs^+^ (2m M, Figure 6A) nor ZD7288 (30 µM, Figure 6B) notably changed the RMPs of DACs in the adult retina. Because ZD7288 and Cs^+^ had similar effects, we pooled their data as shown in Figure 6C. Averaged data shows no significant changes in RMP before (−70 ± 2.08 mV) and during (−70.92 ± 2.04 mV) a blocker (P>0.05, n = 12). However, in the majority of DACs, we found that Cs^+^ (Figure 5A) and ZD7288 (Figure 6B) reduced the frequency of spontaneous activity in DACs, respectively. To quantify the changes in the spontaneous firing frequency, we measured the firing rate (in Hz) before and during the application of an HCN channel blocker. To obtain the percentage change, we first subtracted the firing rate before the drug from the firing rate during the drug. We then divided that difference by the firing rate before the drug and finally multiplied the result by 100. Excitation and inhibition by a drug are defined by more than a 10% increase and less than a 10% decrease, respectively. The percentage in between was considered no change. We found that an inhibition of spontaneous activity frequency was observed in 7 out of 11 DACs (Figure 6D, yellow bar). Two DACs showed no effect of the blockers on spontaneous activity frequency (Figure 6D, orange bar). In addition, HCN inhibitors excited two DACs tested (Figure 6D, red bar).

**Figure 6.**
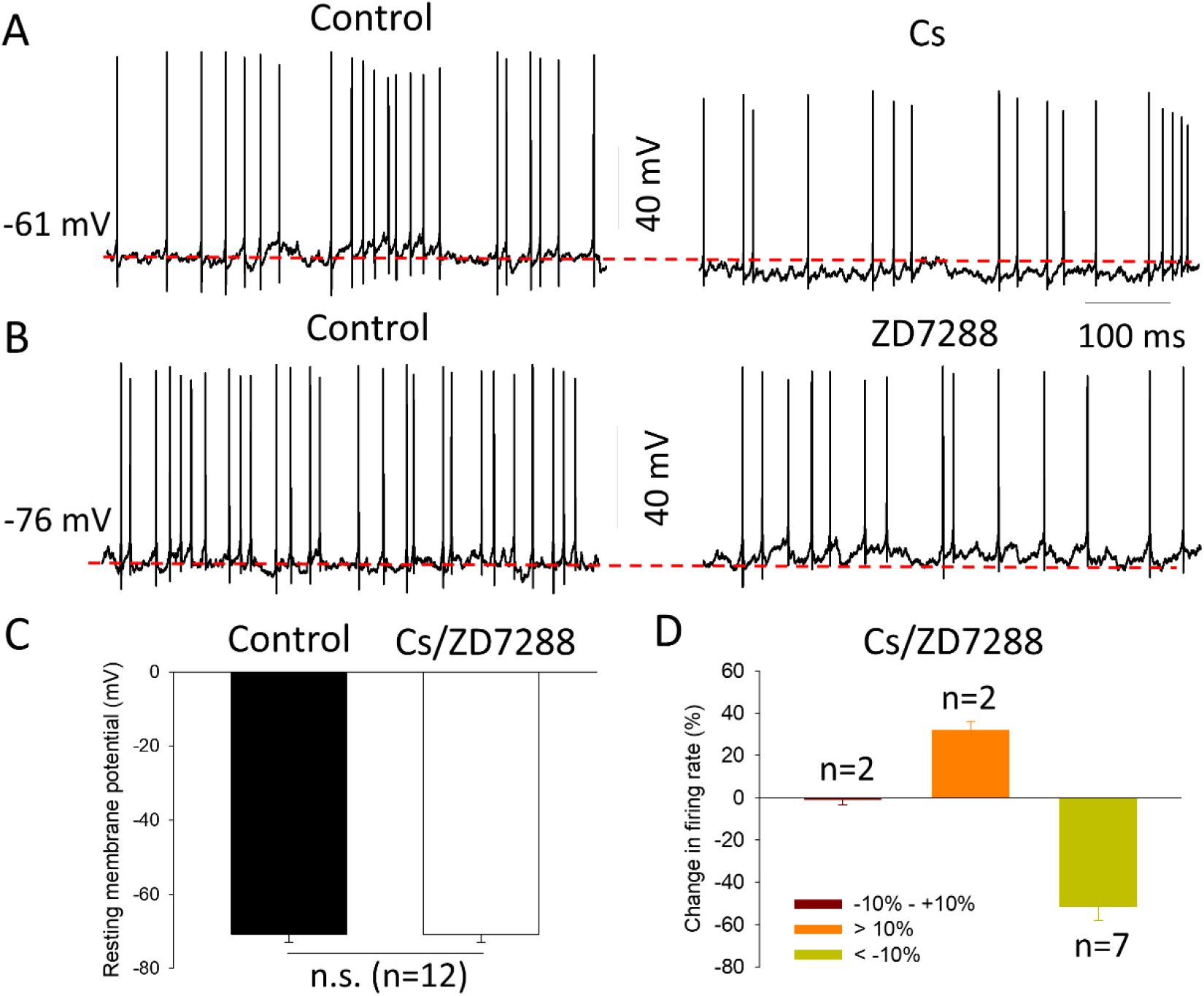
Effect of HCN channel blockers on the spontaneous firing activity of DACs. Whole-cell current-clamp recordings were made on RFP-marked DACs in flat-mount retinas as described in Figure 4. 2 mM Cs **(A**) and 30 µM ZD7288 (**B**) reduced the frequency of spontaneous activity of DACs, respectively, but neither of them notably changed their resting membrane potential (also see **C**). In a total of 11 cells tested (**D**), Cs or ZD7288 excited 2 DACs (red bar, firing rate change>10%), inhibited 7 DACs (yellow bar, firing rate change <-10%), and did not affect on 2 DACs (orange bar, fire rate change between-10% and +10%)). See text for details. Dashed red lines in **A** and **B** indicate the level of the resting membrane potential.

Although varying effects of HCN channel blockers were found, the blockade of HCN channels suppresses the spontaneous activity in the majority of DACs.

## Discussion

We have demonstrated for the first time the expression and distribution of HCN1, HCN2, and HCN4 throughout the retina during development and have described the expression and function of HCN channels in DACs. Specifically, our results clearly demonstrate that the expression of HCN1, HCN2, and HCN4 isoforms begins as early as P4, indicating that HCN channels appear in retinal neurons before the onset of visual experience. Interestingly, HCN1 was expressed in rods/cones, amacrine cells, and retinal ganglion cells (RGCs) at P4, but later in bipolar cells by P12. Differentially, HCN2 and HCN4 were each expressed in amacrine cells and RGCs at P4, but later in bipolar cells by P8 and in later rods/cones by P12. The results suggest that during early development, photoreceptors, and bipolar cells use different isoforms to form HCN channels. More interestingly, these differences disappear by P12, the day around eye-opening. The pattern of HCN1, HCN2, and HCN4 expression in P12 was similar to that in adults, indicating that by P12, HCN channels appear to be fully developed in retinal neurons before eye-opening. This allows them ready to regulate visual information processing after eye-opening^10, 12, 19^. In addition, although we do not identify HCN-expressing retinal neurons other than DACs, the results support the previous study about the contribution of HCN channels in cone bipolar cells to the formation of glutamatergic waves that is critical from the visual circuit development^20, 26^. The results also provide a possibility that M2 and M4 ipRGCs, two of six ipRGC subtypes, express HCN channels that mediate their phototransduction during development and impact visual circuit development^22, 24, 28^.

Our results further demonstrate that HCN1, HCN2, and HCN4 were expressed in approximately half of DACs at P12. However, the expression of each isoform was extended to almost all of DACs in the adult retina. One cause of this difference between ages could be due to the light-induced expression of HCN proteins after eye-opening in adult DACs. Another possible interpretation could be that the expression of HCN channel isoforms is age-dependent in DACs. This assumption is supported by age-dependent upregulation of HCN channels in other tissues^37, 38^. Regardless, adult DACs contain every HCN channel isoform tested, suggesting that they are capable of forming functional channels.

Our electrophysiology data demonstrate that HCN channel isoforms do form heterogeneous functional channels in DACs with a current step-evoked membrane potential sag. The hyperpolarization-evoked potential sag is pharmacologically confirmed to be mediated by HCN channels. The results reveal a specific subtype of amacrine cells that express HCN channels, which extends the previously reported HCN channel-expressing amacrine cells^16, 23, 25^. It is worth noting that although DACs contain the three HCN channel isoforms tested at a protein level, we observed a small percentage of no-sag-expressing DACs. The lack of voltage sag in these neurons may be caused by space-clamp problems because DACs have relatively large somata and dendritic fields in a flat-mount retina^29^.

HCN channels can set and limit the RMPs of a neuron in the central nervous system by allowing the K^+^/Na^+^ influx with a permeability ratio range from 3:1 to 5:1. We found no significant changes in the RMP of DACs while HCN channels were pharmacologically blocked. Although the RMP remains unchanged, the blockade of HCN channels altered the frequency of the spontaneous firing activity. The decrease in the frequency was observed in the majority of DACs, which is consistent with the results obtained in a subpopulation of brain dopamine neurons^39-41^. In agreement with other studies on brain dopamine neurons^42, 43^, the blockade of HCN channels either did not change or increased the frequency of the spontaneous firing activity in a minority of DACs. Therefore, the role of HCN in controlling spontaneous firing is rather complicated. This complication in the retina may be caused by a light/dark-adapted state while the recording is performed due to short exposure of fluorescence light to the retina for the visualization of DACs. In addition, since rods, cones, and bipolar cells are upstream neurons of DACs^44, 45^, our results do not rule out the possibility that HCN channel blockers act on these neurons and then indirectly influence the spontaneous firing activity of DACs. Future studies will address this alternative hypothesis. Overall, the regulation of the spontaneous firing activity by HCN channels in DACs indicates that in addition to previously reported persistent sodium channels^8^, HCN channels may play a critical role in generating spontaneous activity that is associated with tonic dopamine release in the developing and mature retina.

## Acknowledgements

The authors thank Elizabeth Alessio and Laura Gunther for their generous assistance. This work was supported by the Oakland University Provost’s Undergraduate Student Research Award and NIH/NEI Grant R21EY031405, R01EY033808 and R15EY034305.

## Disclosure

**E.J. Romano**, None; **D.-Q. Zhang**, None.

## Notes

### Competing Interest Statement

The authors have declared no competing interest.

